# Architecture of the cardiac transverse-axial tubular system across different mammalian species

**DOI:** 10.64898/2026.01.10.698376

**Authors:** Joachim Greiner, Frédéric Sonak, Wesley Dean Jones, Josef Madl, Kathrine Albertine Ryeng, Igor R Efimov, Zafar Iqbal, Thomas Seidel, Peter Kohl, Eva A. Rog-Zielinska

**Affiliations:** Institute for Experimental Cardiovascular Medicine, University Heart Center Freiburg · Bad Krozingen, and Faculty of Medicine, Albert-Ludwig University of Freiburg, Germany; CIBSS Centre for Integrative Biological Signalling Studies, Albert-Ludwig University of Freiburg, Germany; Institute of Marine Research, Tromsø, Norway; Department of Biomedical Engineering, Northwestern University, Chicago, IL, USA; Institute of Cellular and Molecular Physiology, Friedrich-Alexander-Universität Erlangen-Nürnberg, Erlangen, Germany

**Keywords:** Transverse-Axial Tubular System (TATS), T-tubules, Cardiomyocytes, Confocal Microscopy, Inter-Species Differences, Comparative Physiology

## Abstract

Cardiac excitation–contraction coupling relies on a pancellular network of regular cardiomyocyte surface membrane invaginations, termed the transverse-axial tubular system (TATS). The TATS is ubiquitously present in adult mammalian cardiomyocytes, enabling efficient structural and functional coupling of sarcolemma and intracellular Ca^2+^ stores. However, TATS ultrastructural characteristics across species, and their relation to cardiomyocyte morphology and physiological parameters such as heart rate, remain largely unexplored.

Here, we quantified TATS and cardiomyocyte features in a large confocal microscopy dataset (78 3D volumes) obtained from tissue slices across eight species (mouse, rat, rabbit, pig, horse, elephant, whale, and human). We developed and applied a semi-automated image analysis pipeline to quantify mean cytosolic distances to the nearest TATS (Cyto-TATS_min_, a measure inversely related to TATS density), transverse-to-axial tubule ratio, and cardiomyocyte dimensions. Cyto-TATS_min_ and transverse tubule fraction differed substantially between species, with the lowest Cyto-TATS_min_ in mouse and highest in human. Within species, except mouse, rat, and horse, Cyto-TATS_min_ was positively correlated with cardiomyocyte cross-sectional area. Across all species, Cyto-TATS_min_ correlated with species’ life span and body weight, and was inversely correlated with average resting heart rate.

Our findings reveal structural scaling principles within species differences in cardiac cellular ultrastructure and provide a resource for studying TATS organisation in health and disease. As TATS remodelling is a common hallmark of cardiac pathology, awareness of species-differences is needed to guide the design and interpretation of translational research.

## Introduction

Cardiac contraction arises from well-coordinated shortening of individual cardiomyocytes. Within each working cardiomyocyte, near-synchronous activation of contractile units is initiated by a sharp rise in cytosolic Ca^2+^. The fast and spatially harmonised Ca^2+^ increase is enabled by an elaborate intracellular membrane system, the transverse–axial tubular system (TATS), which plays a crucial role in excitation–contraction coupling.

The TATS is a complex polymorphic network of tubular surface membrane invaginations, present in adult mammalian striated muscle cells, such as cardiomyocytes. Its membrane is continuous with the surface sarcolemma, and its luminal content with the bulk extracellular fluid. Close structural and functional coupling of TATS and the sarcoplasmic reticulum, the cells’ main Ca^2+^ store, enables the rapid and synchronised rise in cytosolic Ca^2+^ levels and – ultimately – uniform and efficient contraction of the whole heart.

While the TATS in atrial cells appears to be less regular, containing a higher proportion of axial elements,^1,2^ the mature TATS in healthy ventricular cardiomyocytes displays a high order of intrinsic spatial organisation, with transverse elements near Z-lines displaying a ‘striation’-like appearance.^3^ TATS structure is remarkably preserved across cardiomyocytes within a species; however, notable differences have been mentioned between species. These inter-species differences include the diameter of individual tubules (ranging from 20 to 600 nm, with larger species reported to have wider TATS),^4,5^ fraction of total plasmalemma contained within TATS (estimated to be between 20% in guinea pig and rabbit, and 60% in mouse and rat), fraction of cardiomyocyte volume occupied by TATS (estimated to be between 1% in guinea pig and 4% in mouse and rat), TATS density (understood to be higher in smaller animals such as rodents, where TATS appears more mesh-like), and fraction of transverse elements (thought to be higher in larger mammals).^6–18^ The architecture of TATS mouth regions also displays species differences − heavily convoluted in mouse, yet fairly open and unobstructed in rat, rabbit, pig, and human.^4^ It has been hypothesised previously that the complexity and density of the TATS also correlates with cell size (e.g. larger cells were suggested to contain a more extensive network), or heart rate (in smaller species with higher heart rates the TATS is thought to be more complex).^19,20^ That said, most of the prior reports did not systematically, directly, or quantitatively evaluate species differences in TATS structure. It is unknown how differences in experimental protocols (e.g. influence of different sample preparation protocols, microscopy modality, use of structural imaging vs. electrophysiological measurements, nature of quantitative analysis) would affect meta-study-based conclusions. Furthermore, many reports focused on qualitative, rather than quantitative assessments.

Deeper knowledge of differences in TATS structure across species, and of the correlations between TATS structure and functional parameters, would help to better understand TATS function in health and disease. The latter is especially important because the TATS has been shown to undergo substantial remodelling in disease, including changes in density and diameter, loss of regularity, or changes in orientation of individual tubules relative to main cell axis.^3,19,21–23^ Furthermore, as the majority of experimental cardiac research relies on the use of animal models, robust quantitative data on the structure of TATS would aid more informed selection of appropriate models to study cardiac function in health and disease, and ease the interpretation and translation of data.

Here, we provide a comparative, quantitative, confocal microscopy-based analysis of TATS structure in cardiac ventricular tissue from commonly used laboratory species and other large mammals – mice, rats, rabbits, pigs, elephant, horse, whale, and human. We further examine how TATS structure and cardiomyocyte morphology correlate with typical physiological parameters for the various species – resting heart rate, body size, and life span.

## Methods

### Ethical Approvals & Animal Handling

All data were sourced with ethical approval from the institutions providing tissue access. Investigations included samples irrespective of animal sex. Investigations on mice (*Mus musculus*), rats (*Rattus norvegicus*), rabbits (*Oryctolagus cuniculus*), and pigs (*Sus scrofa domesticus*) conformed to German animal welfare laws (TierSchG and TierSchVersV) and were compatible with the guidelines stated in Directive 2010/63/EU of the European Parliament on the protection of animals used for scientific purposes. They were approved by local Institutional Animal Care and Use Committees in Germany (Regierungspräsidium Freiburg, X-16/10R, TVA G17–160, X19/01R, and X-20/05R), unless stated otherwise. Laboratory animals were kept in a 12-hour light/dark cycle at room temperature with food and water *ad libitum.* Horses (*Equus caballus*) and elephants (*Elephas maximus*), from which tissue was obtained, belonged to the regular diagnostic pool of the Institute of Veterinary Pathology at the Justus-Liebig-University Giessen, and were either euthanised in accordance with animal welfare regulations (due to illnesses) or had died from natural causes. Whale tissue from minke whales (*Balaenoptera acutorostrata*) was obtained from the Institute of Marine Research, Tromsø (these animals were sacrificed irrespective of subsequent use of heart tissues in the current study). None of the horse, elephant, or whale samples came from an animal used in animal experiments. Transmural cardiac tissue samples from healthy human hearts were obtained through Northwestern University, Chicago, IL, USA from donor hearts rejected for clinical transplantation, where the donors passed away due to non-cardiac reasons. De-identified tissue was provided by the organ procurement organization Gift of Hope (Chicago, IL, USA), and the protocol has been approved by the Northwestern University Institutional Review Board (IRB).

Out of a total of 78 3D cardiomyocyte image volumes, 24 have previously been published as part of the development of CM segmentation methodology.^24^ Tissue sources included in this study are listed in Table 1.

**Table 1:**
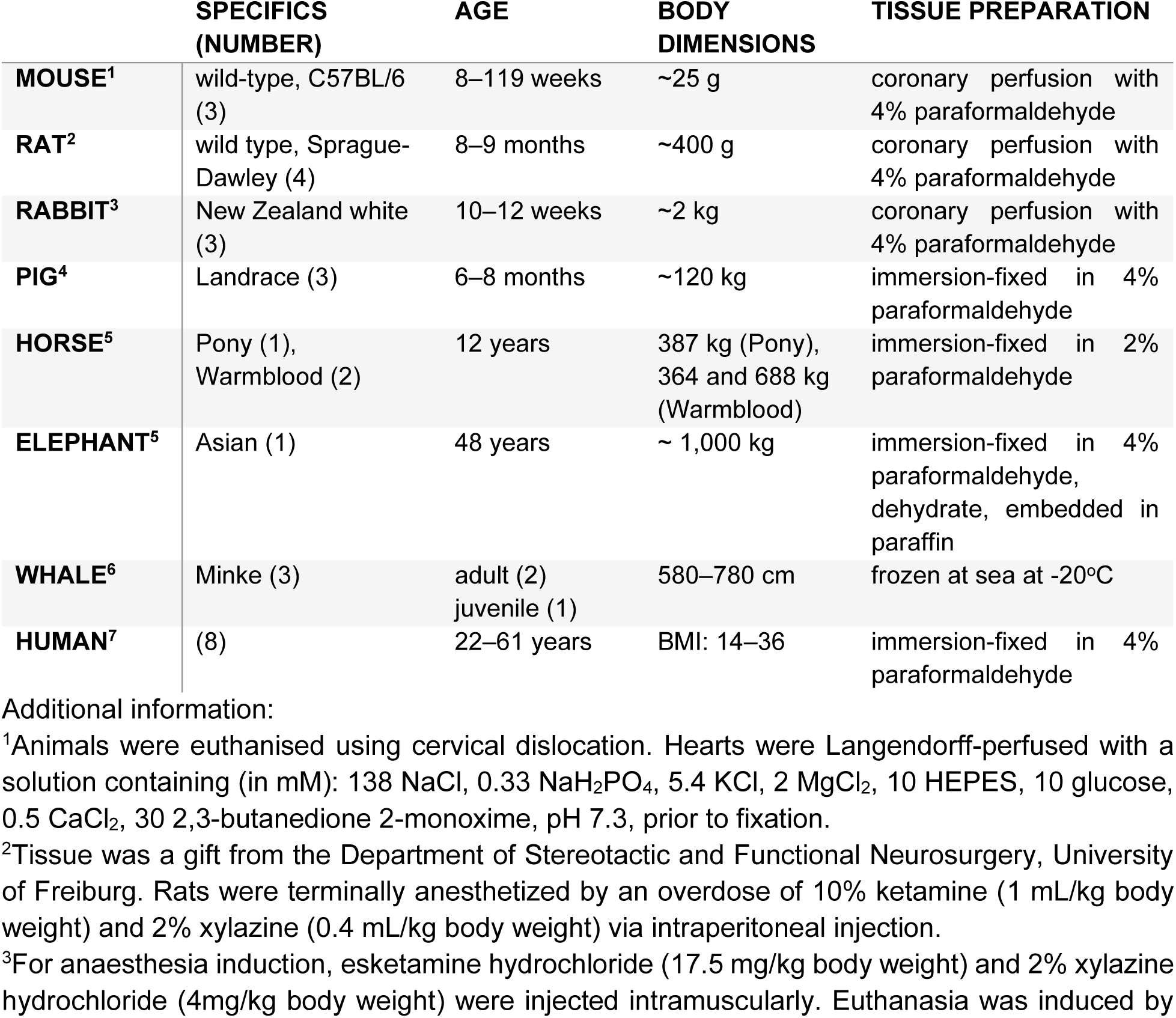
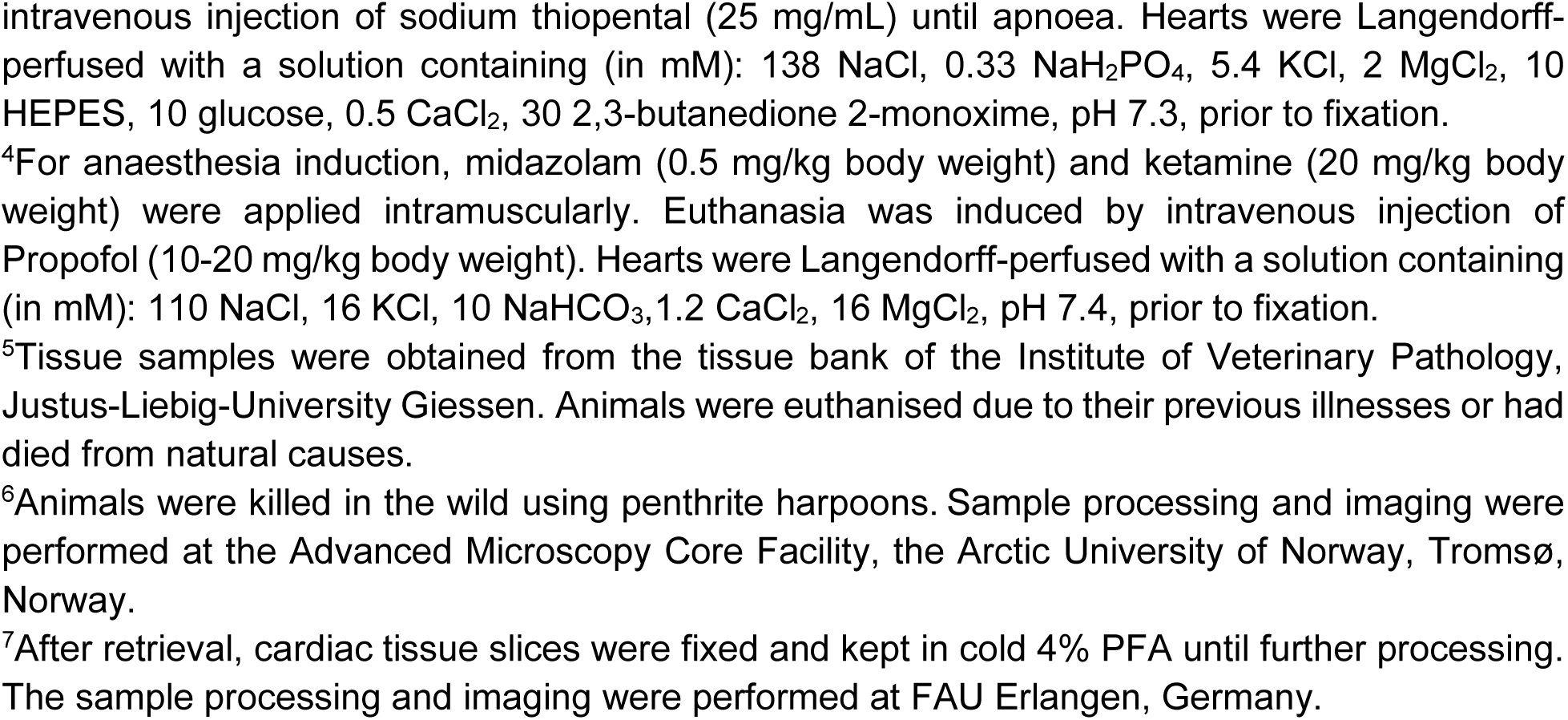
Sources of tissue used in this study.

|  | **SPECIFICS**<br>**(NUMBER)** | **AGE** | **BODY**<br>**DIMENSIONS** | **TISSUE PREPARATION** |
| --- | --- | --- | --- | --- |
| **MOUSE<sup>1</sup>** | wild-type, C57BL/6 (3) | 8–119 weeks | ~25 g | coronary perfusion with 4% paraformaldehyde |
| **RAT<sup>2</sup>** | wild type, Sprague-Dawley (4) | 8–9 months | ~400 g | coronary perfusion with 4% paraformaldehyde |
| **RABBIT<sup>3</sup>** | New Zealand white (3) | 10–12 weeks | ~2 kg | coronary perfusion with 4% paraformaldehyde |
| **PIG<sup>4</sup>** | Landrace (3) | 6–8 months | ~120 kg | immersion-fixed in 4% paraformaldehyde |
| **HORSE<sup>5</sup>** | Pony (1), Warmblood (2) | 12 years | 387 kg (Pony), 364 and 688 kg (Warmblood) | immersion-fixed in 2% paraformaldehyde |
| **ELEPHANT<sup>5</sup>** | Asian (1) | 48 years | ~ 1,000 kg | immersion-fixed in 4% paraformaldehyde, dehydrate, embedded in paraffin |
| **WHALE<sup>6</sup>** | Minke (3) | adult (2)<br>juvenile (1) | 580–780 cm | frozen at sea at -20°C |
| **HUMAN<sup>7</sup>** | (8) | 22–61 years | BMI: 14–36 | immersion-fixed in 4% paraformaldehyde |

Additional information:
<sup>1</sup>Animals were euthanised using cervical dislocation. Hearts were Langendorff-perfused with a solution containing (in mM): 138 NaCl, 0.33 NaH<sub>2</sub>PO<sub>4</sub>, 5.4 KCl, 2 MgCl<sub>2</sub>, 10 HEPES, 10 glucose, 0.5 CaCl<sub>2</sub>, 30 2,3-butanedione 2-monoxime, pH 7.3, prior to fixation.
<sup>2</sup>Tissue was a gift from the Department of Stereotactic and Functional Neurosurgery, University of Freiburg. Rats were terminally anesthetized by an overdose of 10% ketamine (1 mL/kg body weight) and 2% xylazine (0.4 mL/kg body weight) via intraperitoneal injection.
<sup>3</sup>For anaesthesia induction, esketamine hydrochloride (17.5 mg/kg body weight) and 2% xylazine hydrochloride (4mg/kg body weight) were injected intramuscularly. Euthanasia was induced by
intravenous injection of sodium thiopental (25 mg/mL) until apnoea. Hearts were Langendorff-perfused with a solution containing (in mM): 138 NaCl, 0.33 NaH<sub>2</sub>PO<sub>4</sub>, 5.4 KCl, 2 MgCl<sub>2</sub>, 10 HEPES, 10 glucose, 0.5 CaCl<sub>2</sub>, 30 2,3-butanedione 2-monoxime, pH 7.3, prior to fixation.
<sup>4</sup>For anaesthesia induction, midazolam (0.5 mg/kg body weight) and ketamine (20 mg/kg body weight) were applied intramuscularly. Euthanasia was induced by intravenous injection of Propofol (10-20 mg/kg body weight). Hearts were Langendorff-perfused with a solution containing (in mM): 110 NaCl, 16 KCl, 10 NaHCO<sub>3</sub>, 1.2 CaCl<sub>2</sub>, 16 MgCl<sub>2</sub>, pH 7.4, prior to fixation.
<sup>5</sup>Tissue samples were obtained from the tissue bank of the Institute of Veterinary Pathology, Justus-Liebig-University Giessen. Animals were euthanised due to their previous illnesses or had died from natural causes.
<sup>6</sup>Animals were killed in the wild using penthrith harpoons. Sample processing and imaging were performed at the Advanced Microscopy Core Facility, the Arctic University of Norway, Tromsø, Norway.
<sup>7</sup>After retrieval, cardiac tissue slices were fixed and kept in cold 4% PFA until further processing. The sample processing and imaging were performed at FAU Erlangen, Germany.

### Sample Preparation and Imaging

For most non-human samples, chemically-fixed left ventricular tissue was cut into transmural fragments with a cross-sectional area of ∼0.8 cm × 0.8 cm (apart from elephant, where tissue was provided in form of paraffin-embedded 80 µm sections; and whale, where tissue had been frozen by placing it in a freezer at -20°C; see Table 1). Fixed tissue was embedded in 4 % low melting agarose (Carl Roth, Karlsruhe, Germany), glued to the sample stage of a vibratome (7000 smz, Campden Instruments Ltd., Loughborough, UK) with a cyanoacrylate glue, and cut into 50–300 μm thick sections (cut using blade displacements of 1.5 mm amplitude at a frequency of 60 Hz, with an advance speed of 0.05 mm/s). Slices were carefully freed from any remaining agarose using forceps and washed with phosphate-buffered saline (PBS; in mM: 137 NaCl, 2.7 KCl, 8.1 Na_2_HPO_4_, 1.5 KH_2_PO_4_; pH 7.4). Paraffin-embedded elephant tissue slices were dewaxed, rehydrated using graded concentrations of ethanol, and washed in PBS. Whale tissue was cryo-sectioned, immersion-fixed in 2% paraformaldehyde, and washed with PBS. The samples were incubated with 0.5 % Triton-X100 for 30 minutes at room temperature on an orbital shaker. The tissue was washed three times using PBS supplemented with 0.01% Tween-20. Slices were incubated with wheat germ agglutinin (WGA) conjugated to CF488 (10–40 μg/mL, Biotium, Hayward, CA, USA) on a custom-built orbital shaker at 37 °C for up to 48 hours. After washing three times in PBS, samples were mounted on cover glasses with FluoroMount G (SouthernBiotech, Birmingham, AL, US) using a compression-free mounting approach.^25^ The samples were cured in an environmentally-controlled chamber at 42 % humidity (using oversaturated potassium carbonate solution) for 1–3 days before sealing with clear nail polish.

Fixed left-ventricular human samples were washed thoroughly with PBS and incubated for a minimum of 16 h in PBS containing 30% sucrose. Afterwards, the tissue was embedded in Tissue-Tek O.C.T compound (Sakura, Torrance, CA, USA) and cooled to -20 °C for 20 min before cutting into 100 µm-thick sections using a cryostat (CM3050, Leica, Wetzlar, Germany). Slices were washed with PBS twice and permeabilised overnight at 4 °C in PBS containing 1% bovine serum albumin, 5 % normal goat serum, and 0.25 % TritonX-100. Afterwards, slices were washed with PBS three times and stained with AlexaFluor647-conjugated WGA (40 µg/mL, Thermo Fisher, Rheinfelden, Germany) and DAPI (40 µg/mL, Sigma, Darmstadt, Germany) in PBS for a minimum of 6 h in the dark. Stained slices were embedded in microscopy mounting medium (Fluoromount G, SouthernBiotech, Birmingham, AL, US) on microscope slides, covered with coverslips (#1.5), and cured for a minimum of two days at 40–50% relative humidity. Mouse, rat, rabbit, pig, horse, and elephant tissue was imaged at the SCI-MED facility of the Institute for Experimental Cardiovascular Medicine, Freiburg University, using an inverted laser scanning confocal microscope (Leica TCS SP8 X, Leica Microsystems, Wetzlar, Germany) with a glycerol objective (HC PL APO CS2 63× NA 1.4; Leica Microsystems). Laser power was linearly raised during imaging at increasing depth to reduce scattering-induced signal intensity attenuation. Tile scanning was used to increase the area of observation. Whale tissue was imaged at the University of Tromsø using an inverted laser scanning confocal microscope (Zeiss LSM 880, Zeiss, Oberkochen, Germany) with an oil objective (Plan-Apochromat 63× NA 1.4; Zeiss). To acquire 3D volumes of non-human species, 1,024 × 1,024 pixel 2D images with pixel sizes of (90–180 nm)^2^ were recorded as a z-stack series with 90–180 nm z-spacing. To match voxel sampling, image axes with native sampling smaller than 120 nm were downsampled via mean averaging (kernel size 2).

Human tissue was imaged at FAU Erlangen using laser scanning confocal microscopy (Zeiss LSM 780, Zeiss) with an oil objective (Plan-Apochromat 63× NA 1.4; Zeiss). 1,280 × 1,280 pixel 2D images with pixel sizes of (96 nm)^2^ were recorded as a z-stack series with 200 nm z-spacing. Laser power was increased with depth to compensate for signal attenuation. To match voxel sampling, x- and y-image spacing was downsampled using mean averaging (kernel size 2). Our datasets are provided for open-access and include 3D left ventricular cardiomyocyte data from: mouse (7 stacks), rat (7 stacks), rabbit (6 stacks), pig (8 stacks), horse (12 stacks), elephant (4 stacks), whale (10 stacks), and human (24 stacks). Average stack dimensions are 364 × 188 × 54 µm^3^ (non-human), and 123 × 123 × 25 µm^3^ (human).

### Cardiomyocyte Segmentation and Analysis

We segmented cardiomyocyte instances in 3D using a deep learning-based image analysis framework as described previously.^24^ First, we trained and applied a 3D U-Net for content-aware image restoration (CARE)^26^ to reduce spatially varying blur and noise in WGA-labelled image volumes. Next, we segmented individual cardiomyocytes using the DTWS-MC method and interactively proofread the resulting masks in SegmentPuzzler.^24^ For morphometric analyses, we conducted a principal component analysis of voxel coordinates of each cell mask and used the resulting vectors to define a local coordinate system. We used three orthogonal eigenvectors to span the coordinate axes, with the vectors sorted by decreasing eigenvalue, and the primary cell axis corresponding to the largest eigenvalue. Cell length, width, and depth were defined as the mask extent along the first, second, and third axes, respectively. Cross-sectional area was measured in a plane orthogonal to the first axis (spanned by the second and third axes). We analysed the 30 largest cardiomyocyte masks in each image volume of non-human tissues, and the 5 largest cardiomyocyte masks in the image stacks of human tissue. These cut-offs were set *a priori* to balance coverage and truncation of cardiomyocyte masks, because stacks obtained using non-human samples contained substantially larger volumes than those from human tissue. We report per-cell average values for cross-sectional area and cell width and depth. We do not report average cardiomyocyte length because most image stacks were too small to contain sufficient numbers of complete (non-truncated by volume boundaries) cardiomyocytes. In addition, acquisition depth differed across species.

### TATS Segmentation and Analysis

We applied image processing for each cardiomyocyte to compute the cytoplasm-to-nearest-TATS voxel distance (Cyto-TATS_min_, an inverse measure of effective TATS density) and to extract the transverse-to-axial tubule fraction. First, we filtered structures with a low spatial frequency (assumed to be non-TATS) with a per-slice white 2D top-hat filter (box structuring element with radius of 5 voxels) using clEsperanto.^27^ We thresholded WGA by computing Li’s minimum cross-entropy threshold^28^ with scikit-image,^29^ multiplied by an empirically determined factor of 0.75. The threshold was calculated using voxels constrained within the cardiomyocyte mask. For the Cyto-TATS_min_ analysis, we used a single volume-wide threshold to generate 3D distance maps using the SignedMaurerDistanceMap function in ITK.^29^ For TATS orientation analysis, thresholds were calculated independently for every 2D slice. To exclude non-TATS structures, we eroded the cardiomyocyte mask in-plane by 6 voxels and intersected the thresholded WGA with it. We then computed a TATS skeleton and transformed it into a local coordinate system aligned with the cardiomyocyte’s long and short axes, computed with a PCA from the corresponding 2D mask (axes ordered by decreasing eigenvalues). Next, we classified each skeleton voxel using a 3 x 3 in-plane neighbourhood centred on that voxel: axial, if there were two neighbouring voxels forming a segment parallel to the long axis of the cell; transverse, if there were two neighbouring voxels forming a segment parallel to the short axis of the cell, or other (e.g. diagonal structures). To classify transverse elements cut orthogonally to their long axis (appearing as elliptical elements in 2D sections) correctly, despite not meeting the neighbourhood criterion, we further classified skeleton voxels as transverse if a difference-of-Gaussians filter (standard deviations of 2 and 4 voxels) matched their location.

### Visualisation and Statistics

Data points shown represent the mean value across all analysed cardiomyocytes within a single stack, and stacks were the unit of analysis for group comparisons. Overlaid summary statistics show mean ± standard deviation across stacks. For statistical testing across species, we used the Python package pingouin^30^ and a one-way ANOVA on per-stack means, with post hoc comparisons using the Holm procedure to correct for multiple comparisons. A p-value of less than 0.05 was taken to indicate a statistically significant difference between means. For within-species analyses, cells were the unit of analysis, and linear regressions were fitted with ordinary least squares using SciPy.^31^ We used a two-sided Wald t-test to assess whether the fitted slope differed from 0. Two-sided 95% confidence intervals for the fitted mean were computed from the same model in statsmodels.^32^ Power law relations *y* = *ax*^*b*^were fitted analogously to the linear fit, but with the input data being log-transformed (log *y* = log(*a*) + b · log(*x*)). Two-sided 95% confidence intervals for the fitted mean were computed in log space, then back-transformed and adjusted using Duan’s smearing factor calculated from the log-residuals of each fit. Pearson’s r and *p* values were taken from the log-log regression.

## Results

We curated a large dataset of 3D confocal image stacks of cardiac left ventricular tissue labelled using WGA. Part of this dataset was used by us previously to develop tools for quantitative cardiomyocyte segmentation.^24^ Our dataset includes small rodents such as mice and rats; medium-sized lagomorphs such as rabbits; large animals, such as pigs and horses; very large animals such as elephants and whales; and humans. We provide open-source access to 78 3D image stacks, accompanied by a partial or complete instance segmentation of cardiomyocytes and TATS (see Table 1 for species distribution). The protocol for preparing and acquiring images was similar across most species, with deviations when unavoidable, such as handling pre-embedded tissue (elephant), chunk-frozen tissue (whale), or using samples acquired in the process of another project (human).

### The architecture of the TATS network varies between species

We segmented and classified TATS elements with our automated image-processing workflow across eight species (Fig. 1A). Quantitative results are summarised in Table 2; analyses pertain to the following number of cardiomyocytes/stacks/animals: mouse: 210/7/3, rat: 210/7/4, rabbit: 180/6/3, pig: 240/8/3, horse: 357/12/3, elephant: 115/4/1, whale: 300/10/3, human: 120/24/8.

**Figure 1:**
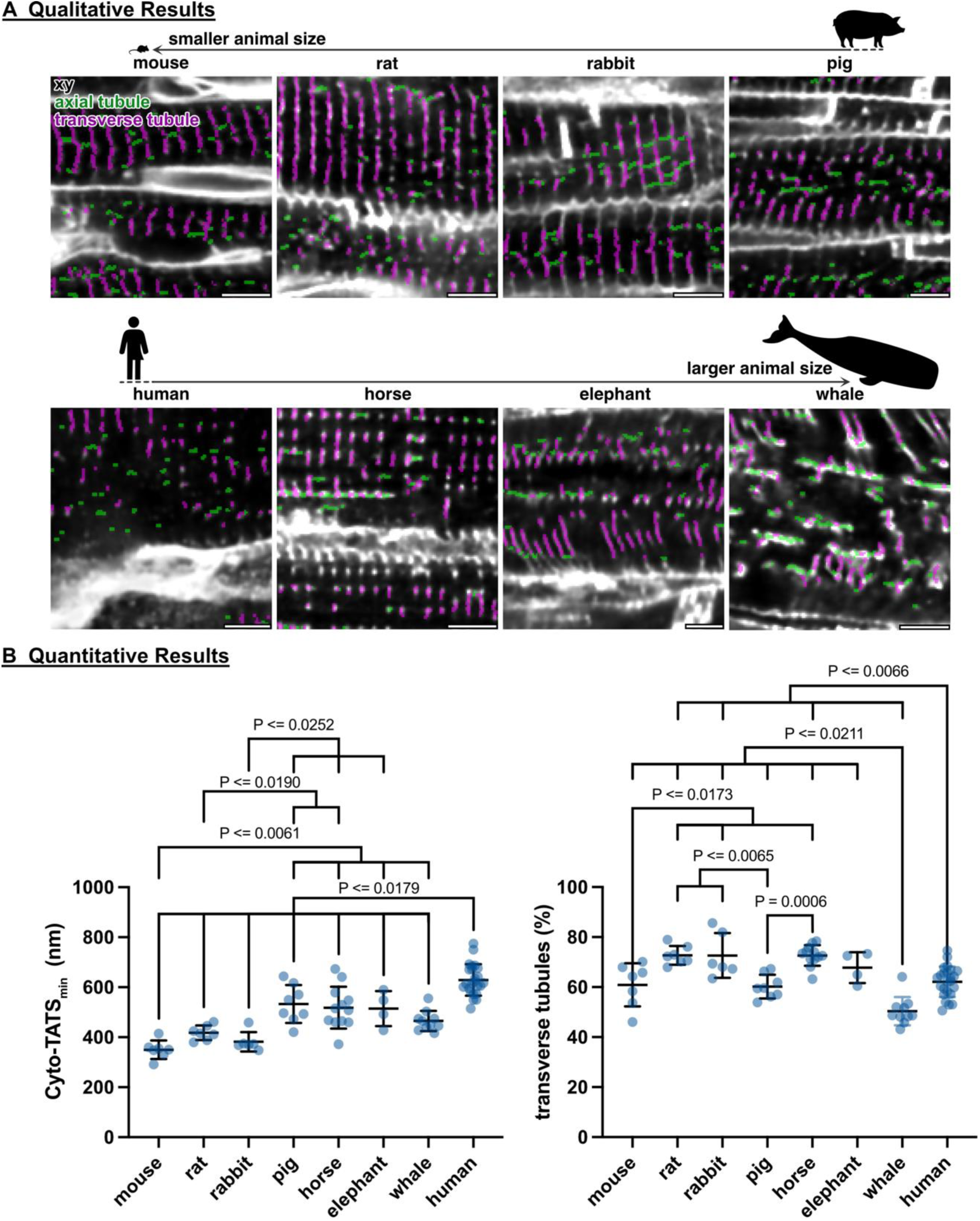
Analysis of left ventricular TATS structure across a range of mammalian hearts. **A:** Representative TATS skeletonisation, derived from 3D confocal microscopy image stacks of WGA-stained tissue slices. Magenta indicates transverse, and green axial tubular elements of the TATS; scale bars: 5 µm. **B:** Quantitative analysis of TATS features: Cyto-TATS_min_, and transverse tubule fraction; each dot represents an average value across all cardiomyocytes of per image stack. Number of cardiomyocytes/stacks/animals here and in all subsequent figures: 210/7/3 (mouse), 210/7/4 (rat), 180/6/3 (rabbit), 240/8/3 (pig), 357/12/3 (horse), 115/4/1 (elephant), 300/10/3 (whale), 120/24/8 (human).

**Table 2:**
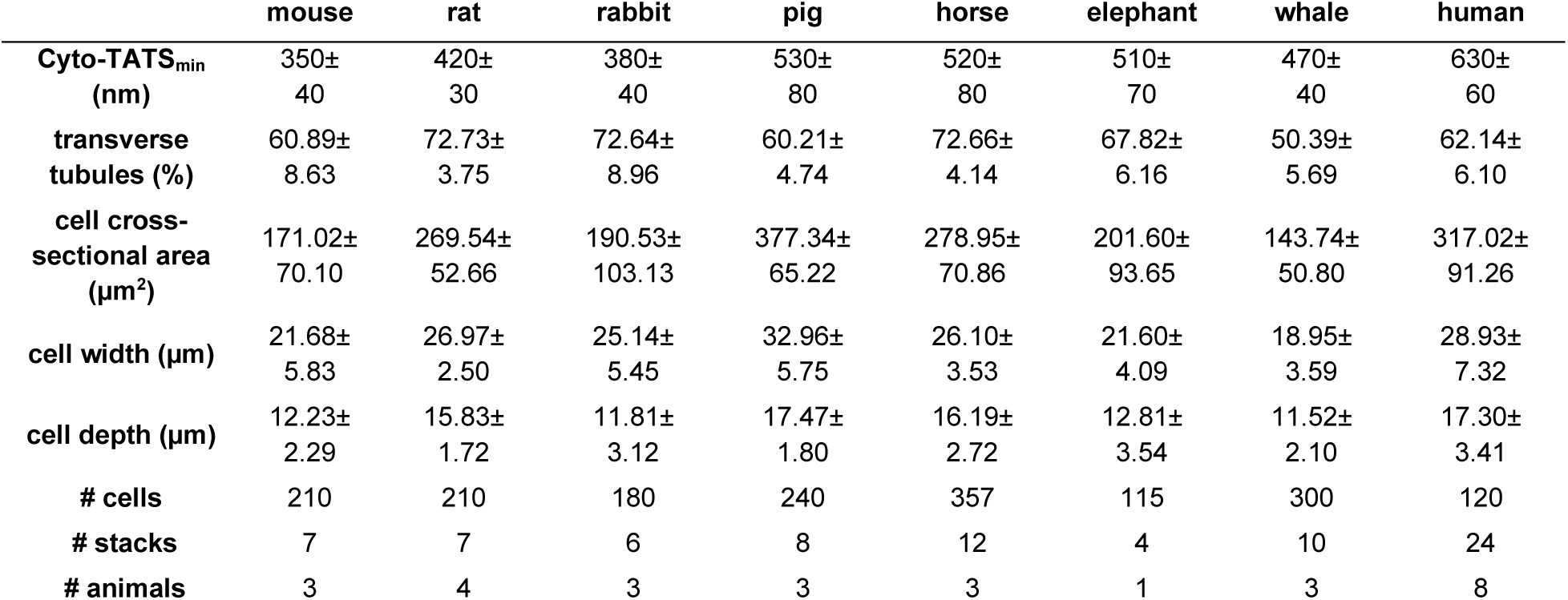
Cardiomyocyte and TATS characteristics of the curated dataset. Data are presented as mean ± standard deviation.

We first assessed the Cyto-TATS_min_ (Fig. 1B left panel). Human cardiomyocytes show significantly highest Cyto-TATS_min_ (630±60 nm; *p* ≤ 0.0179) of all species examined, while Cyto-TATS_min_ was lowest in mouse (350±40 nm) – significantly lower than in pig, horse, elephant, or whale (530±80 nm, 520±80 nm, 510±70 nm, 470±40 nm, respectively; *p* ≤ 0.0061). Rat TATS Cyto-TATS_min_ was significantly lower than pig and horse (420±30 nm vs 530±80 nm and 520±80 nm, respectively; *p* ≤ 0.0190). Rabbit Cyto-TATS_min_ was significantly lower than pig, horse, and elephant (380±40 nm vs 530±80 nm, 520±80 nm, 510±70 nm, respectively; *p* ≤ 0.0252).

Furthermore, we assessed the percentage of transversely oriented tubules in all species (Fig. 1B right panel). Of the species examined, transverse tubule fraction was lowest in whale (50.39±5.69 %), and was significantly lower than all other (i.e. mouse, rat, rabbit, pig, horse, elephant, and human; 60.89±8.63 %, 72.73±3.75 %, 72.64±8.96 %, 60.21±4.74 %, 72.66±4.14 %, 67.82±6.16 %, 62.14±6.10%, respectively *p* ≤ 0.0211). Human cardiomyocytes had a significantly lower transverse tubule fraction than rat, rabbit, and horse (62.14±6.10 % vs 72.73±3.75 %, 72.64±8.96 %, 72.66±4.14 %, respectively; *p* ≤ 0.0066). Mouse and pig cardiomyocytes had a significantly lower transverse tubule fraction compared to rat, rabbit, and horse (60.89±8.63 %, 60.21±4.74 % vs 72.73±3.75 %, 72.64±8.96 %, 72.66±4.14 %, respectively; *p* ≤ 0.0173).

### Correlations between the architecture of TATS and cardiomyocyte dimensions

Examination of the cardiomyocyte dimensions (cross-sectional area, width, and depth; Fig. 2A) revealed species-dependent differences in cardiomyocyte size (Fig. 2B, Table 2). Mouse and rabbit cardiomyocyte cross-sectional areas were significantly smaller than those in pig and human (171.02±70.10 µm^2^, 190.53±103.13 µm^2^ vs 377.34±65.22 µm^2^, 317.02±91.26 µm^2^, respectively; *p* ≤ 0.0149). Whale cardiomyocyte cross-sectional areas were significantly smaller than rat, pig, horse, and human (143.74±50.8 µm^2^ vs 269.54±52.66 µm^2^, 377.34±65.22 µm^2^, 278.95±70.86 µm^2^, 317.02±91.26 µm^2^, respectively; *p* ≤ 0.0333). Elephant cardiomyocyte cross-sectional area was significantly smaller than pig (201.6±93.65 µm^2^ vs 377.34±65.22 µm^2^, *p* = 0.0102).

**Figure 2:**
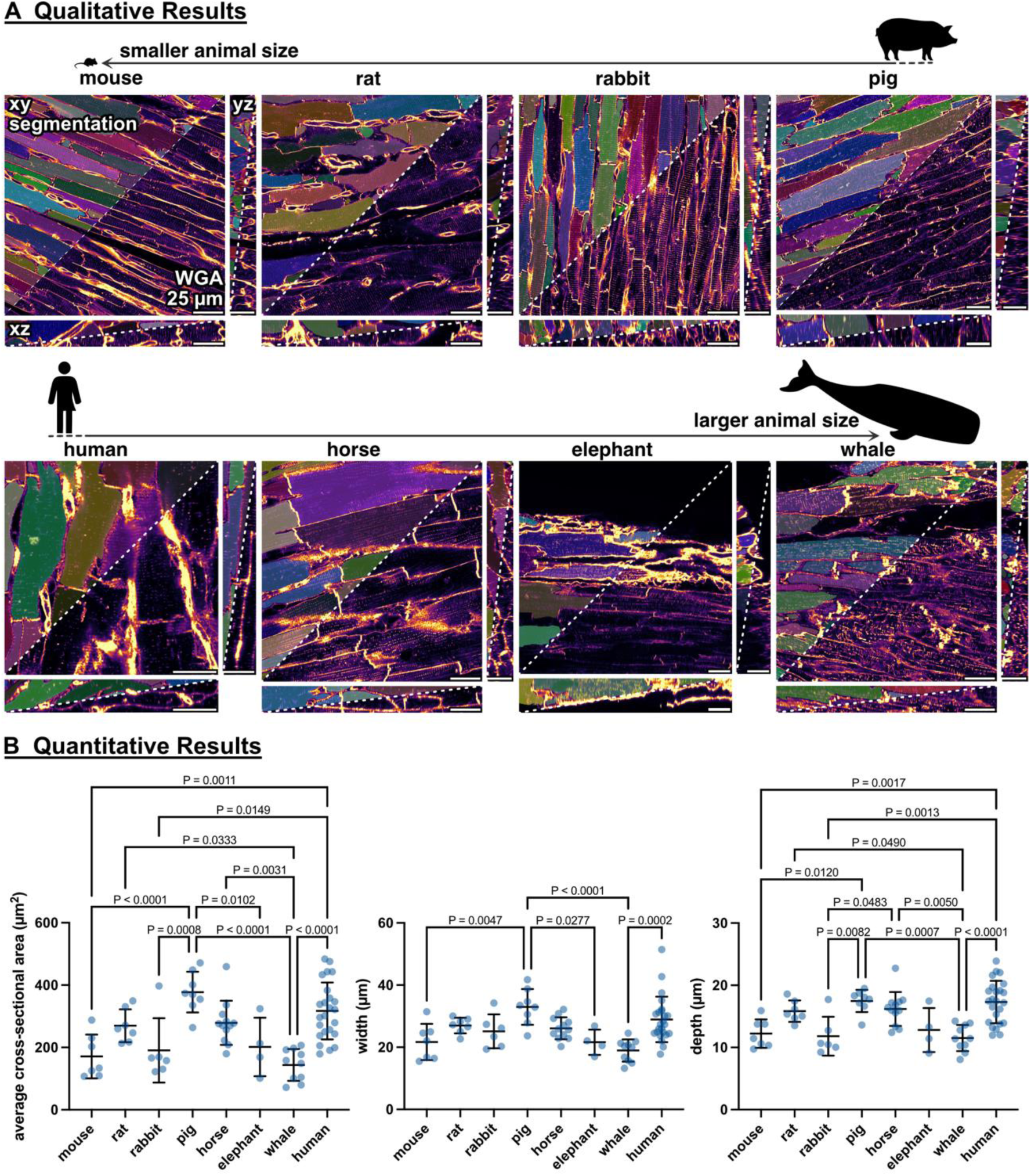
Analysis of left ventricular 3D cardiomyocyte morphology across a range of mammalian hearts. **A:** Representative xy/xz/yz views of WGA-labelled myocardium and cardiomyocyte segmentations. Scale bars: 25 µm. **B:** Quantitative analysis of cardiomyocyte morphology, featuring cross-sectional area, width (2^nd^ smallest dimension), and depth (smallest dimension); each dot represents an average value across all cardiomyocytes per single image stack.

Cardiomyocyte cell width was significantly greater in pig compared to mouse, elephant, and whale (32.96±5.75 µm vs 21.68±5.83 µm, 21.60±4.09 µm, 18.95±3.59 µm, respectively; *p* ≤ 0.0277), and significantly less in whale compared to human (18.95±3.59 µm vs 28.93±7.32 µm; *p* < 0.001).

Finally, cardiomyocyte cell depth was significantly greater in pig and human when compared to mouse, rabbit, and whale (17.47±1.80 µm, 17.30±3.41 µm vs 12.23±2.29 µm, 11.81±3.12 µm, 11.52±2.10 µm, respectively; *p* ≤ 0.012), significantly greater in horse when compared to rabbit and whale (16.19±2.72 µm vs 11.81±3.12 µm, 11.52±2.10; *p* ≤ 0.0483), and significantly greater in rat when compared to whale (15.83±1.72 µm vs 11.52±2.10 µm, *p* = 0.049)

Analysis of the correlation between TATS architecture and individual left ventricular cardiomyocyte morphology within species (Fig. 3, top panel) revealed that Cyto-TATS_min_ was positively correlated with all measures of cardiomyocyte size (cross-sectional area, width, and depth) in elephant (*p* < 0.001), rabbit (*p* < 0.001), and whale (*p* = 0.00163, *p* = 0.00313, *p* < 0.001). Cyto-TATS_min_ was also positively correlated with cardiomyocyte cross-sectional area and depth in human (*p* = 0.0412, *p* < 0.001), and positively correlated with cardiomyocyte cross-sectional area and width in pig (*p* < 0.001). Cyto-TATS_min_ negatively correlated with cardiomyocyte cross-sectional area and width in mouse (*p* = 0.027 and *p* = 0.00226), and cross-sectional area in rat (*p* = 0.0162).

**Figure 3:**
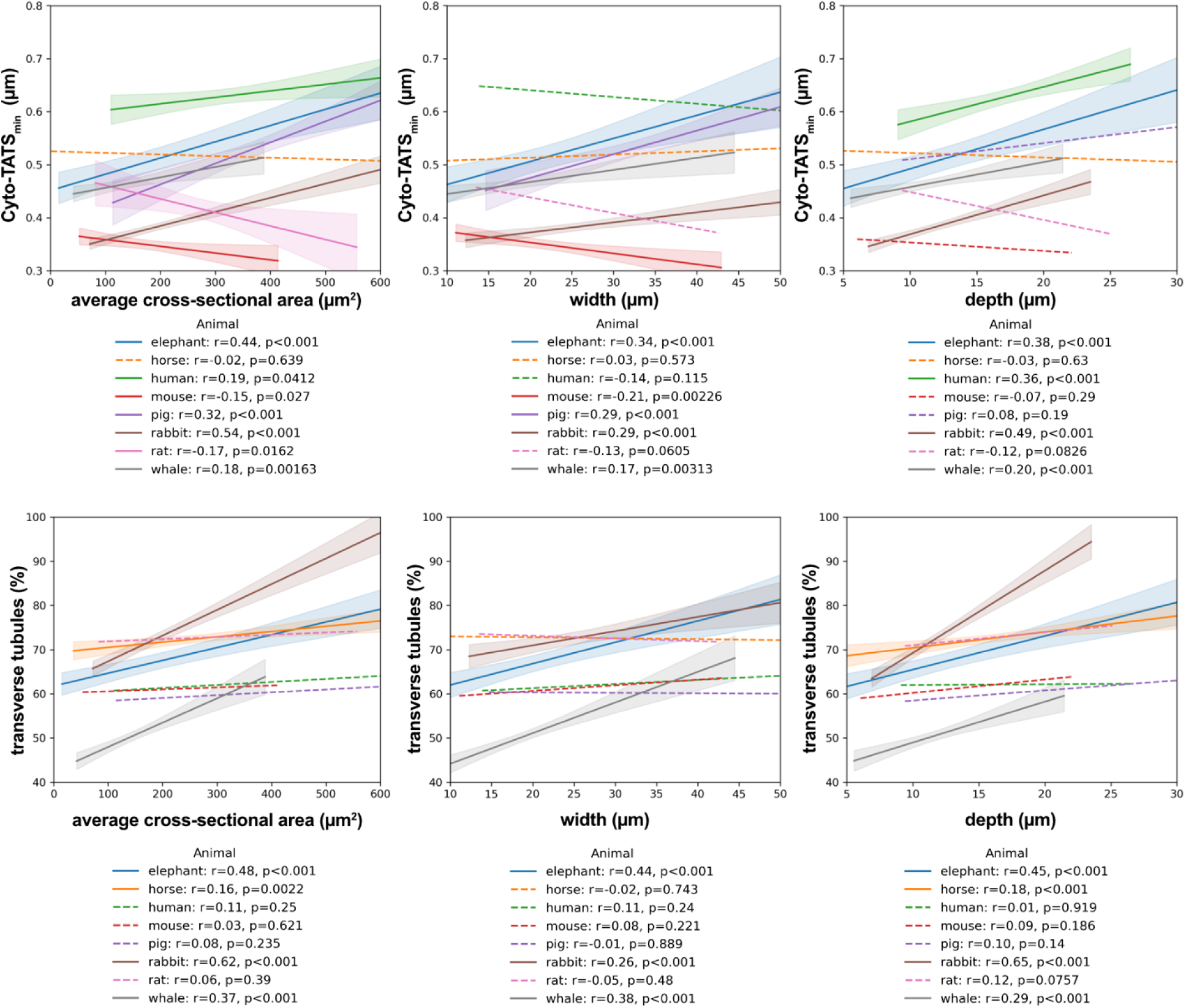
Analysis of the correlations between left ventricular TATS structure and 3D cardiomyocyte morphology across a range of mammalian hearts. The correlations between Cyto-TATS_min_ (top panels, and of the transverse tubule fraction (bottom panels) are shown as a function of cardiomyocyte cross-sectional area (left), width (2^nd^ smallest dimension; middle), and depth (smallest dimension; right). Correlations were analysed using linear regressions. All statistically significant correlations are indicated using a continuous line and include confidence intervals; all statistically non-significant correlations are presented using dashed lines.

Transverse tubule fraction increased with increasing cardiomyocyte cross-sectional area, width, and depth in rabbit (Fig. 3, bottom panel*; p* < 0.001) and elephant (p < 0.001). Transverse tubule fraction increased with increasing cardiomyocyte cross-sectional area and width in whale (*p* < 0.001), and with increasing cardiomyocyte cross-sectional area and depth in horse (*p* = 0.0022, *p* < 0.001).

### Correlation of TATS architecture and physiological parameters

Analysis of the correlation between TATS structure and literature-sourced representative physiological parameters (Fig. 4; and supplementary Table S1) revealed that Cyto-TATS_min_ was higher in species with longer life spans (*p* = 0.0101) and greater body mass (*p* = 0.0481), and lower in animals with higher average resting heart rates (*p* = 0.0426). No correlations were observed between transverse tubule fraction and average life spans, body mass, or average heart rate.

**Figure 4:**
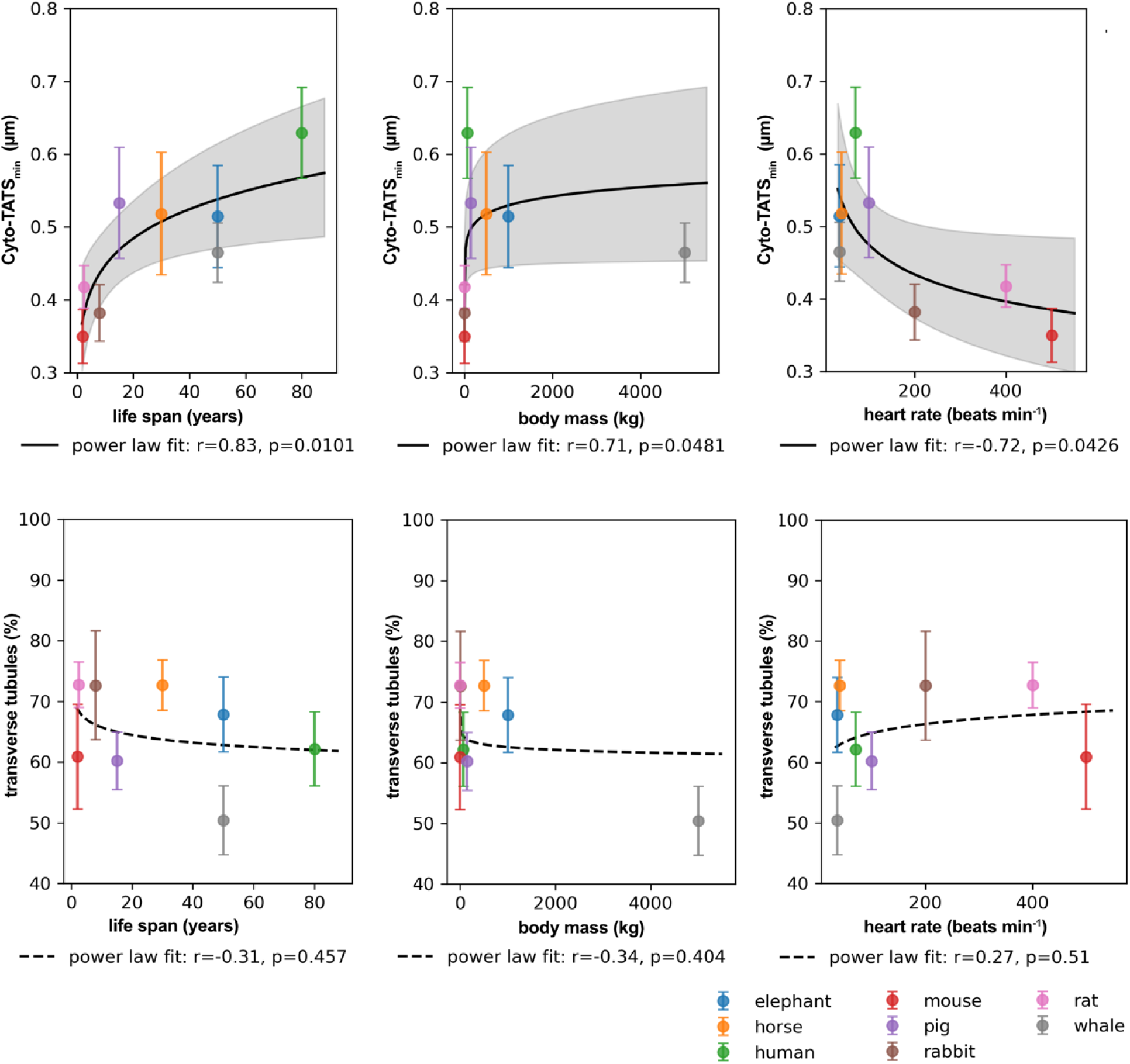
Analysis of the correlations between left ventricular TATS structure and typical physiological parameters across a range of mammalian species. Cyto-TATS_min_ (top panels), and transverse tubule fraction (bottom panels) are shown as a function of average life span (left), body mass (middle), and average resting heart rate (right). Correlations were analysed using nonlinear (power law) regressions, and are provided with confidence intervals.

## Discussion

The TATS is an essential structural and functional component of cardiomyocytes, crucial for efficient and uniform excitation−contraction coupling. Although the importance of the TATS is well recognised, there is limited understanding of how TATS architecture varies across species, despite the widespread use of diverse animal and disease models in research. We characterised the architecture of TATS in left ventricular tissue across multiple species, including common laboratory animals (mouse, rat, rabbit, pig), larger species (horse, elephant, whale), and human; thereby sampling a wide range of physiological parameters such as life span, mass, and heart rate.

Our results indicate that there are clear quantitative differences in TATS density and proportion of transverse tubules (in relation to all TATS elements) across species. TATS density is highest in mouse cardiomyocytes, and lowest in human cardiomyocytes. The comparatively high TATS density in mouse is consistent with previous studies.^9,11,16,33^ The finding that TATS density is lowest in human cardiomyocytes is surprising, as substantially larger mammals were included in our investigation.

TATS density was negatively correlated with a measure of cardiomyocyte size (cross-sectional area, width, or depth) within most of the species examined, apart from mouse, rat (in those two species the correlation was positive), and horse, which suggests that – in contrast to previous suggestions based on atrial cardiomyocytes^2^ – larger cardiomyocytes may not necessarily require a denser TATS network to maintain excitation−contraction synchronicity.

Across all species examined, TATS density was positively correlated with average resting heart rate, and negatively correlated with body mass and lifespan. Of course, lifespan metrics are influenced by extrinsic mortality and sampling context (wild vs captive), and for humans in particular, modern medicine and living conditions will have increased these values to levels that do not fit into a comparison metrics. Furthermore, covariation between body mass and lifespan, both with an inverse association with resting heart rate, is well documented in the literature;^34–37^ thus, these cross-species correlations should be interpreted as associations within a shared allometric scaling framework. That said, there is a plausible link between TATS density and heart rate, as a more extensive coupling of sarcolemma and intracellular Ca^2+^ stores may support more rapid excitation−contraction coupling. Indeed, previous studies have indicated that TATS density correlates with the dynamics of Ca^2+^ transients inside cardiomyocytes.^38–42^

Across the species examined here, transverse tubule fraction was lowest in whale cardiomyocytes. When comparing our results to earlier studies on rodent cardiomyocytes, our results are consistent with a reported transverse tubule fraction of about 60%.^11^ Currently, the exact functional importance of axial elements in healthy ventricular cardiomyocytes is not known, although computational studies suggested that presence of axial elements can aid equilibration of TATS content along cardiomyocytes via intraluminal diffusion.^43^

Our study has a number of applications, for example informing researchers as to the choice of most appropriate research models that best mimic human anatomy. Most basic cardiac research is done using small rodent models, due to the ease of handling, comparatively low cost, fast reproductive cycles, and abundance of recombinatorial genetic tools. However, there is growing concern among cardiac researchers as to the extent to which mice and rats mimic human myocardial structure and function. Here we provide further evidence that mouse and rat TATS structure is not representative of human ventricular cardiomyocytes. Lagomorphs, such as rabbits (dubbed the largest of the small laboratory species) may offer a more relevant model for translational cardiac research. Larger mammals (such as pigs), can mimic human cardiac TATS structure more closely; these are, however, more costly to implement and they benefit less from pre-existing research experience, reference data, and transgenic model systems. Furthermore, while we examined mostly tissue from nominally healthy animals (with the exception of horse and elephant), follow-up research should also examine TATS structure in diseased tissue from the same species. In cardiac pathologies, TATS networks are known to be remodelled, with changes in density and in orientation of individual elements – these changes may precede and be causally linked to cardiac dysfunction.^3,19,21,44^ It would also be of value to comparatively examine TATS architecture in atrial samples, as well as to investigate regional (e.g. inter-chamber, as well as intra-cellular) variability in TATS structure.^41,45,46^.

Our study suffers from a number of technical limitations. As our data was acquired in different laboratories, with tissue preserved using several protocols (in part without stringent control over the mechanical state of the tissue prior to fixation), some of the observed differences (for example in cell cross-sectional area) may thus be related to sample preparation and processing (if that induced a systematic bias between cell contraction states). Furthermore, we relied on confocal microscopy images of tissue stained with WGA, which itself does not stain membranes, but rather the glycocalyx. This approach may introduce a bias against narrower, more difficult to stain TATS elements. Furthermore, as some of our datasets came from non-research animals that either died at an advanced age or were euthanised, our data is confounded by the lack of age-matching. Also, we have no insight into activity levels of animals prior to tissue sampling, which may have affected TATS architecture,^47^ and for one species (elephant), the number of independent biological repeats is one. Finally, for practical reasons, our analyses of correlations between TATS architecture and physiological parameters have focused on average reported values for the species, rather than actual parameters that would have been observed in the individual donor animals.

In terms of image analysis, we didn’t exclude the space occupied by nuclei (about 1 % of cell volume) from our analysis. While most cardiomyocytes are mono- or bi-nucleated, the number of nuclei in pig cardiomyocytes can by up to 8 or 16 per cell,^48^ which could have affected density measurements in this species. While there is no evidence in the density distribution data to suggest that this was the case (perhaps in part because pigs may not only have the largest number of nuclei, but they also have the largest cardiomyocytes; Table 2), this should be addressed in the future.

In conclusion, our study provides a comprehensive, quantitative analysis of ventricular TATS architecture across a range of mammalian species, linking cellular microstructure to organismal physiology. We demonstrate that TATS density is positively correlated with average resting heart rate, and negatively correlated with average life span and body weight. Beyond this, our dataset and analytical framework offer a resource for selecting experimental models of cardiac physiology and pathology. By establishing a reference for TATS organisation in healthy myocardium, we provide groundwork for future studies examining disease-related remodelling of the TATS.

## Acknowledgements

We would like to thank Simone Nübling, Stefanie Perez-Feliz, Cinthia Walz, Pia Iaconniani, Patricia Aparecida Morais Costa, Thomas Kok, Jana Ebeling, and Callum Zgierski-Johnston for technical assistance and advice. We acknowledge the SCI-MED imaging facility at the Institute for Experimental Cardiovascular Medicine in Freiburg for access to the confocal microscope. This research was supported by the German Research Foundation Emmy Noether programme (396913060 to EARZ). JG, WDJ, JM, PK, and EARZ are members of the Collaborative Research Centre SFB1425 of the German Research Foundation (422681845). We thank Kenneth Bowitz Larsen at the Arctic University of Norway, Tromsø, Norway, for assistance with imaging. We thank Yixin Tong, Department of Stereotactic and Functional Neurosurgery, University of Freiburg, for providing rat tissue. We thank the Institute of Veterinary Pathology, Justus-Liebig-University Giessen, for providing horse and elephant tissue.

---

**Supplementary Table 1.**
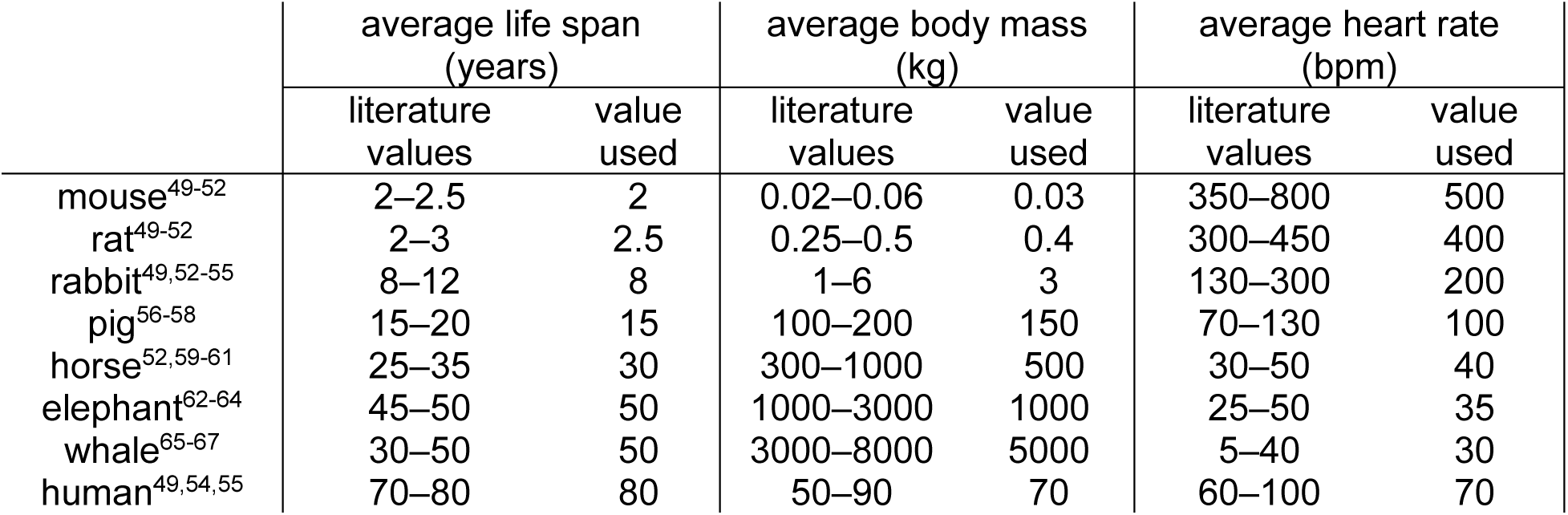
Literature-based typical ranges for species-related life span, body mass and resting heart rate, and values used to assess correlations between TATS structure and physiological parameters in Figure 4.

|  | average life span<br>(years) |  | average body mass<br>(kg) |  | average heart rate<br>(bpm) |  |
| --- | --- | --- | --- | --- | --- | --- |
|  | literature<br>values | value<br>used | literature<br>values | value<br>used | literature<br>values | value<br>used |
| mouse<sup>49-52</sup> | 2–2.5 | 2 | 0.02–0.06 | 0.03 | 350–800 | 500 |
| rat<sup>49-52</sup> | 2–3 | 2.5 | 0.25–0.5 | 0.4 | 300–450 | 400 |
| rabbit<sup>49,52-55</sup> | 8–12 | 8 | 1–6 | 3 | 130–300 | 200 |
| pig<sup>56-58</sup> | 15–20 | 15 | 100–200 | 150 | 70–130 | 100 |
| horse<sup>52,59-61</sup> | 25–35 | 30 | 300–1000 | 500 | 30–50 | 40 |
| elephant<sup>62-64</sup> | 45–50 | 50 | 1000–3000 | 1000 | 25–50 | 35 |
| whale<sup>65-67</sup> | 30–50 | 50 | 3000–8000 | 5000 | 5–40 | 30 |
| human<sup>49,54,55</sup> | 70–80 | 80 | 50–90 | 70 | 60–100 | 70 |

